# Integrated single-cell atlas of human atherosclerotic plaques

**DOI:** 10.1101/2024.09.11.612431

**Authors:** K. Traeuble, M. Munz, J. Pauli, N. Sachs, E. Vafadarnejad, T. Carrillo-Roa, L. Maegdefessel, P. Kastner, M. Heinig

## Abstract

Atherosclerosis, a major cause of cardiovascular diseases, is characterized by the buildup of lipids and chronic inflammation in the arteries, leading to plaque formation and potential rupture. The underlying causal immune mechanisms and alterations in structural cell composition and plasticity driving plaque progression remain incompletely defined. Recent advances in single-cell transcriptomics (scRNA-seq) have provided deeper insights into the roles of immune and non-immune cells in atherosclerosis. However, existing public scRNA-seq datasets often lack comprehensive cell type coverage and consistent annotations, posing challenges for downstream analyses. In this study, we present an integrated single-cell atlas of human atherosclerotic plaques, encompassing 261,747 high-quality annotated cells from carotid, coronary, and femoral arteries. By benchmarking and applying the best-performing data integration method, scPoli, we achieved robust cell type annotations validated by expert consensus and surface protein measurements. This comprehensive atlas enables accurate automatic cell type annotation of new datasets, optimal experimental design, and deconvolution of existing as well as novel bulk RNA-seq data to comprehensively determine cell type proportions in human atherosclerotic lesions. It facilitates future studies by providing an interactive WebUI for easy data annotation and experimental design, while supporting various downstream applications, including integration of genetic association studies and experimental planning.

## Introduction

Atherosclerosis is the primary pathology behind acute ischemic cardiovascular events, like myocardial infarction and stroke^1^. Atherosclerosis is characterized by lipid accumulation and chronic inflammation in the arteries, leading to plaque formation and potential rupture^2,3^. The overarching immune mechanisms and transformation and differentiation processes in vascular cells that reside within the affected arteries, such as vascular smooth muscle cells (SMCs), endothelial cells (ECs), and fibroblasts involved in plaque progression are incompletely understood and under current investigation. The recent advances of single cell transcriptomics (scRNA-seq) gives relevant insights into these processes, unraveling previously undetermined roles of immune and non-immune cells. For example, Wirka *et al*^4^ characterized modulated SMCs that transform into fibroblast-like cells they termed fibromyocytes, which play a protective role in coronary artery disease. Fernandez *et al*^5^ thoroughly investigated the contribution of T cells and macrophages, and identified certain subsets associated with plaque vulnerability.

Many public scRNA-seq datasets of human atherosclerotic plaques are available, but often do not cover the full breadth of disease-contributing cell types, as they are focused on specific subtypes based on cell sorting approaches and preprocessing being applied prior to library preparation and sequencing^6–9^. Additionally, some cell types are easier to harvest in scRNAseq, and hence more abundant in the datasets^10^. Currently, cell type annotations of many of these data sets are not publicly available, making the annotation of cell types and composition of plaques a major challenge that requires expert knowledge. Because most downstream analysis tasks require cell type information, accurate consensus annotation of cell types across datasets is of uttermost importance. Consequently, single cell atlases that integrate and harmonize various published datasets, such as the human lung cell atlas^11^ or the heart cell atlas^12^, emerged as useful references enabling coherent downstream analyses^13^, such as automatic cell type annotation of new datasets^11^, optimal experimental design^14^, interpretation of genetic association studies^11,15,16^ and deconvolution of bulk RNA-seq data sets^17^. Atlases of different tissues are ultimately paving the way towards a comprehensive human cell atlas^18^ that can be used to train foundation models^19–21^, which require vast amounts of data.

To understand atherogenesis and how healthy vascular cells transform into atherosclerotic plaques, first single cell data sets of healthy arteries need to be profiled^22^ to define healthy arterial cell types as a reference. In a second step, it is key to obtain a high-resolution cell type annotation of the cells of atherosclerotic plaques to enable comparisons. First single cell atlases of atherosclerotic lesions^23,24^ have already been proposed. These atlases have several shortcomings, as they are composed mainly of healthy tissue samples, inflating the number of cells in the atlas. As a consequence, the effective number of plaque specific cells is still relatively low, which limits the robustness of cell type annotations and the ability to detect rare but disease-relevant cell types. Moreover, there is currently no evaluation of the consensus of annotations in the field. Finally, existing atlases are limited to two types of arteries, carotids and coronaries, neglecting plaques from other arterial sites like femoral arteries of great relevance to vascular occlusive disease (PVOD). A key practical limitation appears to be the lack of publicly available annotations and model weights to effectively make use of the existing atlases.

For this current study, we have curated an easily accessible plaque cell atlas that encompasses the most comprehensive dataset to date with 261.747 high quality annotated cells from human atherosclerotic carotid, coronary and femoral arteries. We applied the best performing data integration method, selected from a wide spectrum of available models through the most comprehensive model benchmark on a range of metrics specifically evaluated on plaque single cell data sets. The cell type annotations were orthogonally validated with expert annotation consensus and surface protein measurements. We made the annotations and model weights easily accessible by providing an easy-to-use interactive WebUI to automatically annotate new datasets including uncertainty. The performance of the atlas and model was demonstrated and validated in several downstream tasks, such as automated cell type annotation, planning of future experiments with scPower^14^, and combining the vast information of single cell data with the big sample sizes of bulkRNA-seq datasets by deconvolution with BayesPrism^17^. Overall, this comprehensive and robust atlas marks a significant step forward in understanding the complex mechanisms of atherosclerosis and enhances the utility of transcriptomic profiling technologies in cardiovascular research.

## Results

### Integration of public datasets into one atlas

We collected all publicly available single cell datasets of plaque from carotid, coronary and femoral arteries covering a total of 261,747 cells (after quality control (QC)) of diverse cell types and pathologies (see Table 1). Figure 1 shows the workflow to construct the single cell plaque atlas. First, all datasets were pre-processed as described in the original publications. If no detailed description was available, we applied our own dataset-specific quality metrics (see methods section for details). Subsequently, we applied ambient RNA correction^25^ and doublet detection^26^ on each dataset/sample independently.

**Table 1:** All publicly available scRNA-seq datasets for plaque tissues in humans from coronary, femoral and carotid arteries.

| Dataset | Plaque location | # Cells | # Samples | Accession number |
| --- | --- | --- | --- | --- |
| Pan et al.<sup>37</sup> | carotid | 8.867 | 3 | GSE155512 |
| Alsaigh et al.<sup>38</sup> | carotid | 34.716 | 3 | GSE159677 |
| Jaiswal et al.<sup>36</sup> | carotid | 5.567 | 2 | GSE179159 |
| Dib et al.<sup>8</sup> | carotid | 25.194 | 6 | GSE210152 |
| Fernandez et al.<sup>5</sup> | carotid | 9.984 | 7 | GSE224273 |
| Slysz et al.<sup>9</sup> | carotid | 27.540 | 4 | GSE234077 |
| Pauli et al.<sup>35</sup> | carotid | 1.278 | 8 | GSE247238 |
| Bashore et al.<sup>39</sup> | carotid | 77.112 | 18 | GSE253904 |
| Wirka et al.<sup>4</sup> | coronary | 11.756 | 8 | GSE131778 |
| Emoto et al.<sup>6</sup> | coronary | 1.624 | 2 | GSE184073 |
| Chowdhury et al.<sup>7</sup> | coronary | 22.671 | 12 | GSE196943 |
| Slysz et al.<sup>9</sup> | femoral | 35.438 | 9 | GSE234077 |
| **Total** |  | **261.747** | **80** |  |

**Figure 1:**
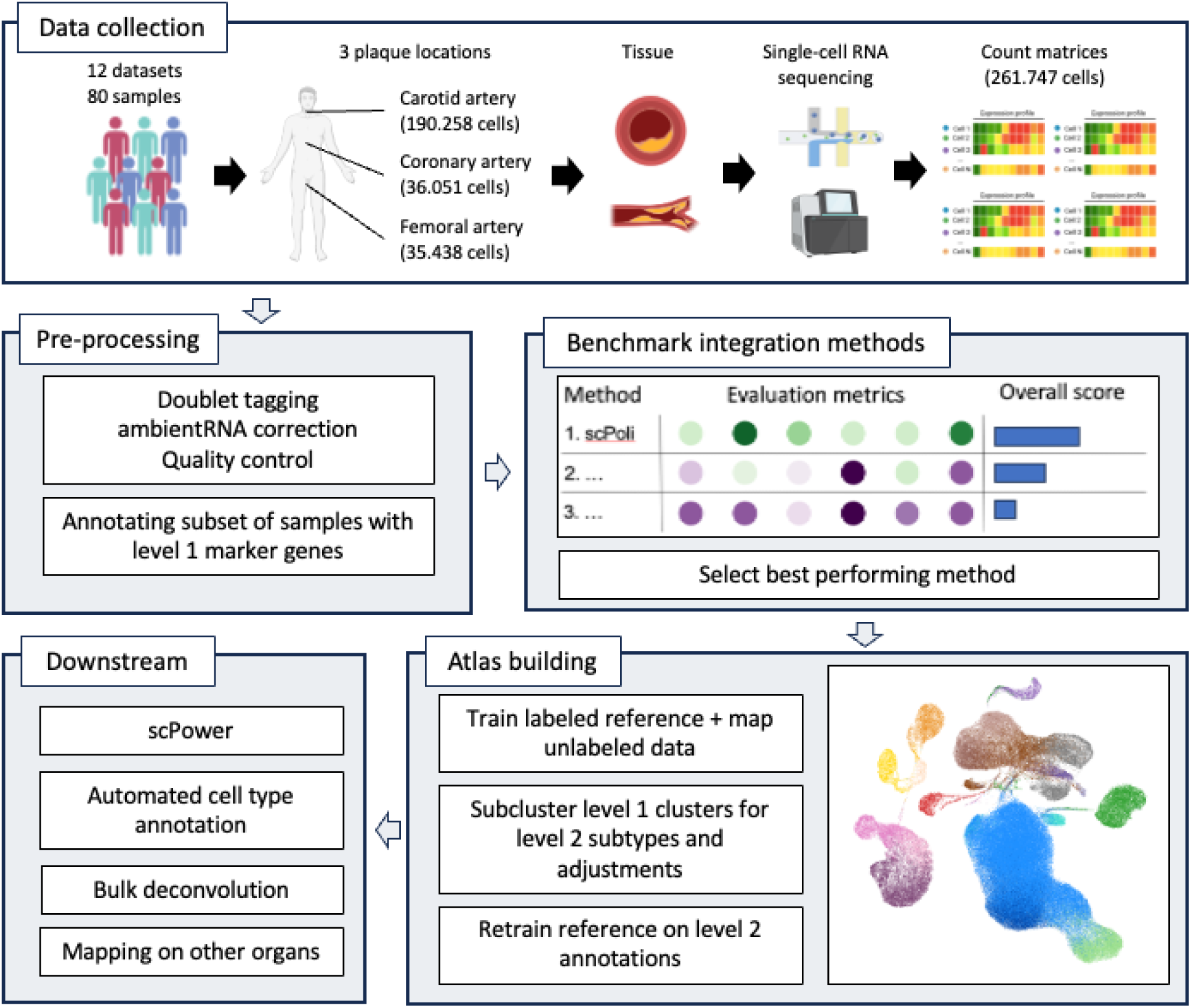
Workflow to construct the single cell plaque atlas. 1) All publicly available scRNA-seq datasets from atherosclerotic plaques from carotid, coronary and femoral arteries were collected from the NCBI GEO database. 2) The same preprocessing pipeline was applied to all datasets. Doublets were tagged, counts were corrected for ambient RNA and cells and genes were filtered as described in the original publication. A subset of samples was manually annotated with level 1 marker genes for the subsequent benchmark of integration methods. 3) A benchmark of integration methods on five batch correction and five biological conservation metrics identified the best performing method, scPoli. 4) ScPoli was applied on the subset of annotated samples to build the atlas. Subsequently sub clustering was applied to major level 1 cell types to identify finer grained level 2 sub cell types. Finally, the atlas was retrained on the level 2 annotations and all cells were assigned to a cell type. 5) The annotated plaque atlas was used in the following downstream applications: designing future experiments with scPower, using the atlas as a reference for automating the cell type annotation and bulk RNA dataset deconvolution.

Next, different datasets were integrated into one latent space to eliminate technical batch effects and conserve biological signals. To select the best method for this task, we performed an extensive benchmark with scbi-metrics^27^ of commonly used integration methods: *scVI*^28^*, Harmony*^29^*, LIGER*^30^*, scANVI*^31^*, scGen*^32^*, scPoli*^33^ and *baseline PCA*. The benchmark was evaluated on several metrics in the categories batch correction and bio conservation, where the latter requires cell type labels. For this reason, we manually annotated a subset of 11 samples using a carefully curated table of plaque-specific marker genes (see Figure 2A). This table, the most comprehensive human sample collection to date, includes genes previously utilized for annotating plaque cells in scRNA-seq studies. The 11 samples were manually annotated incrementally until the total number of cells in each cell type was at least 600 (see Suppl. Figure 1). This enabled us to achieve robust annotations and to cover all major cell types that we labeled as “level 1” annotation. Next, we compared methods based on these annotations and other metrics. As expected, the baseline PCA had the worst batch correction score out of all methods (see Suppl. Figure 2). Surprisingly, scVI performed the worst in bio conservation, while having a high batch correction score, indicating an over correction. Fine tuning the scVI model with the scANVI method demonstrated a substantial improvement in the bio conversation score. Overall, the method scPoli outperformed all other methods in both bio conservation and batch correction metrics and was hence used to integrate all the remaining samples as well.

**Figure 2:**
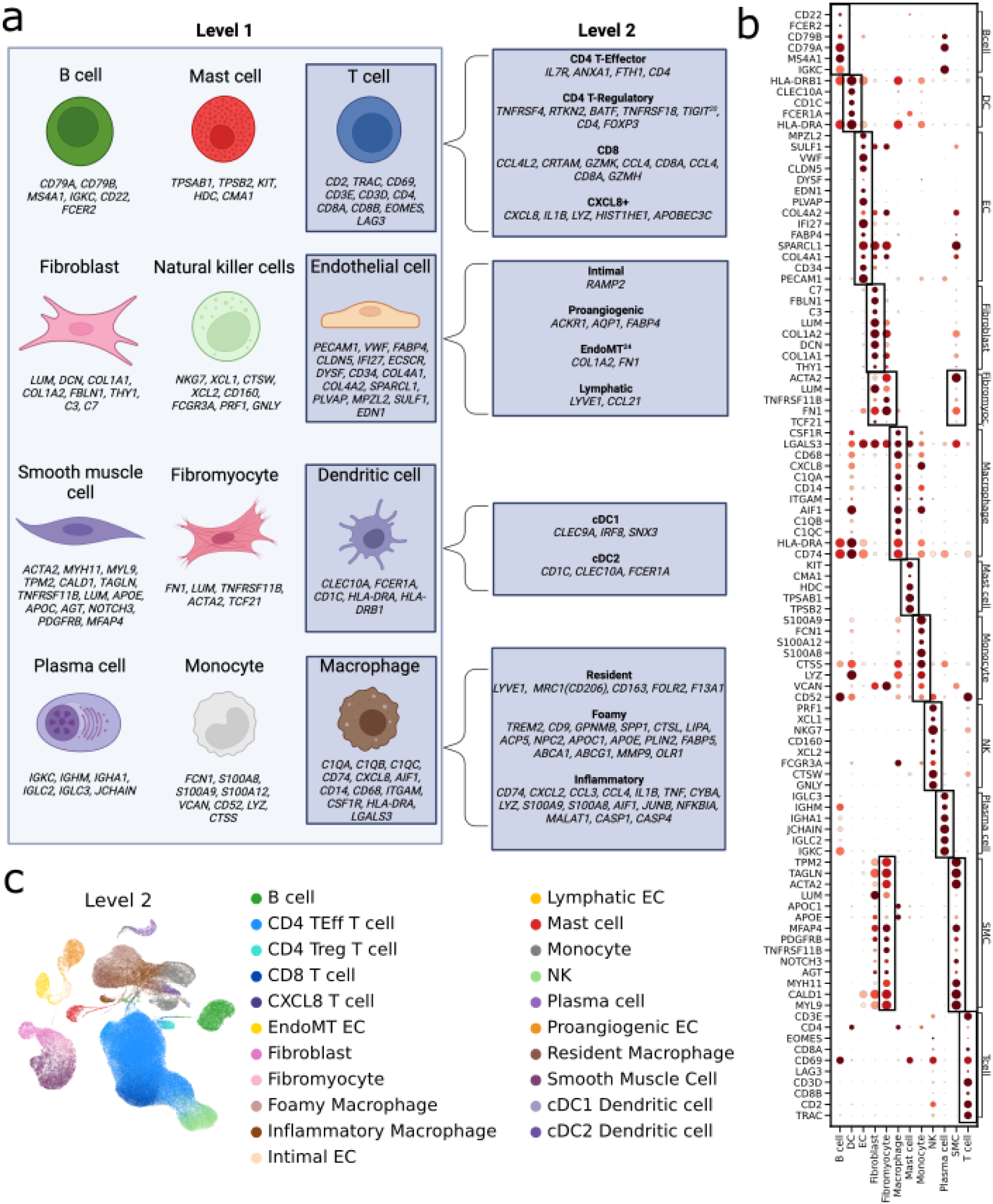
Cell type annotation. Panel A) shows an overview of marker genes in atherosclerotic plaques previously used in scRNA-seq datasets to annotate cells. Level 1 cell types refer to major cell types commonly used in studies, while level 2 refers to a finer grained sublevel annotation of specific cell types with a distinct biological function. Level 1 marker genes for B cells^23,38,40–43^, mast cells^6,40,43–46^, T cells^5,38,43,47^, fibroblasts^23,43,44^, NK cells^8,38^, EC^23,38,40,43,44^, SMC^23,38,40,43,44,48–50^, fibromyocytes^4,23,44^, DC^6,23,40,45,51^, plasma cells^23,44^, monocytes^6,23,40^, macrophages^5,6,38,40,44,52^. Level 2 marker genes for CD4 T-Effector^9^, CD4 T-Regulatory^9^, CD8 T cell^9^, CXCL8+ T cell^9^, intimal EC^23,53^, proangiogenic EC^23,54,55^, EndoMT^23,56^, lymphatic EC^23,57^, cDC1^8^, cDC2^8^, resident macrophages^23,40,58,59^, foamy macrophages^5,23,40,58–65^, inflammatory macrophages^5,40,58^. Panel B) shows a dot plot of the level 1 marker genes in the predicted cell types in the reference atlas. The color coding shows normalized gene expression of the marker genes (y-axis) in each cell type (x-axis), with light red indicating low expression and dark red indicating high expression. Genes are repeated if they are considered marker genes for several cell types. Panel C) shows a UMAP projection of the reference atlas, illustrating level 2 cell types. Each dot represents a cell and colors indicate the level 2 cell type annotation specified in the legend. Created with BioRender.com.

**Figure 3:**
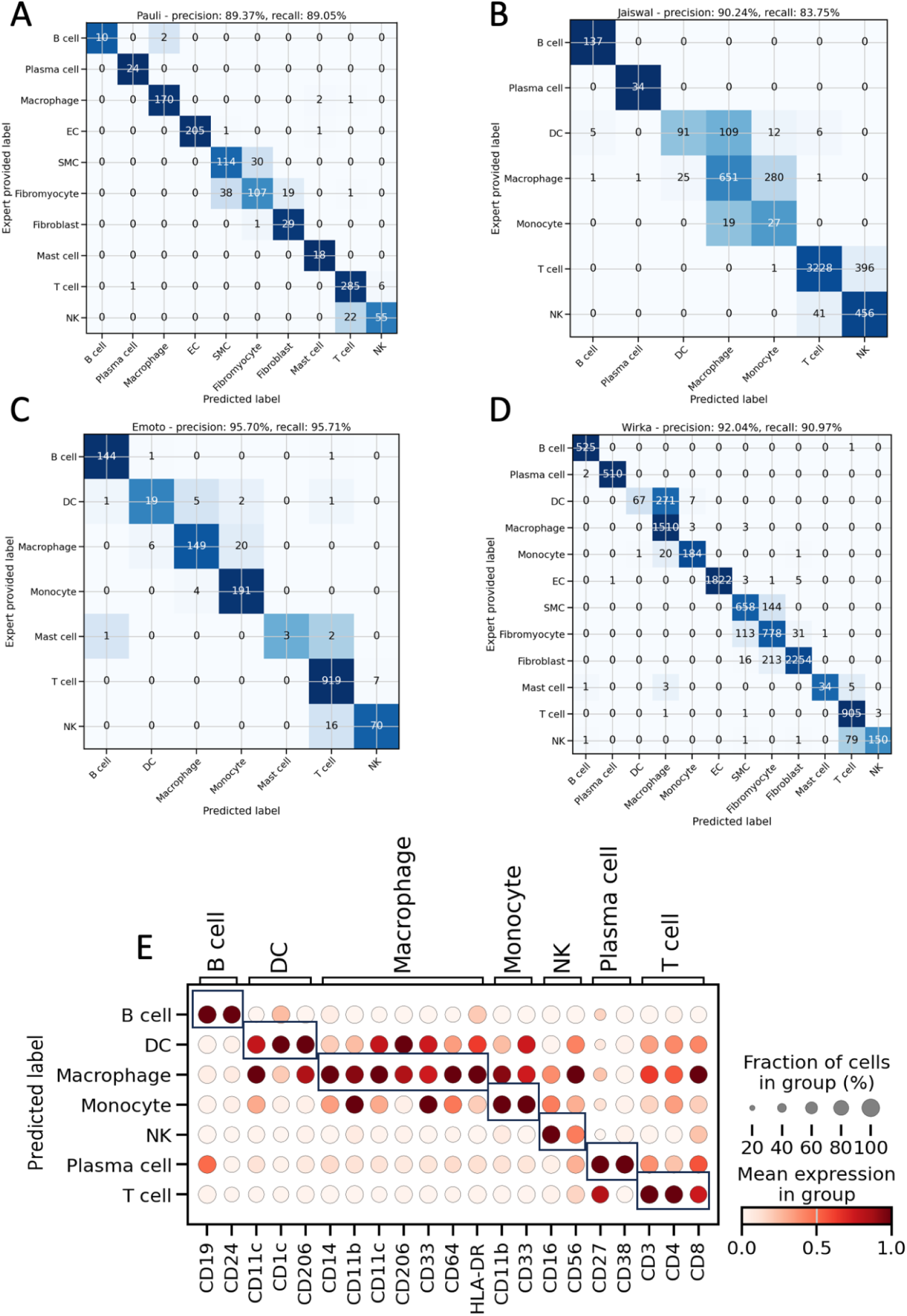
Validation of cell type label transfer. Panels A-D show the confusion matrices for different data sets (data set name given in the panel headers) providing the number of cells with specific combinations of expert annotations (“Expert provided Label” on the y-axis) and labels predicted by the atlas (“Predicted Label” on the x-axis). To compensate for the differences in cell type abundance the colormap is normalized along the rows and the overall precision and recall are weighted according to their abundance. Panel E depicts the dot plot of surface markers grouped according to cell types (x-axis) in a CITE-seq sample to orthogonally validate the cell type annotations of the atlas (y-axis). The highly specific expression pattern of the surface protein is depicted with black boxes. The color coding shows normalized protein expression of the proteins encoded by the marker genes, with light red indicating low expression and dark red indicating high expression.

We then proceeded with the atlas building step by transferring level 1 cell type annotations to the remaining unlabeled cells using the scArches^34^ method. Subsequently, we looked for finer grained cell types, which we refer to as “level 2”, where we focus on cell types that have distinct biological functions that have previously been described in the literature. To achieve this, all cells of a given level 1 cell type were sub-clustered and annotated with curated plaque-specific marker genes for level 2 cell types (see Figure 2A). This implies that we only annotated robust subtypes that were functionally characterized in previous studies. Additionally, we adjusted potentially misannotated subclusters, where we identified for example subclusters expressing T cell specific markers among the cells annotated as natural killer cells (NK) in level 1. These cells were relabeled as T cells. Subclusters expressing macrophage specific marker genes among the cells annotated as dendritic cells (DC) in level 1 could be relabeled as macrophages. Finally, the scPoli model was retrained on the level 2 annotations to produce the data integration model for all downstream applications. The UMAP-projected embedding obtained after level 2 training is depicted in Figure 2C and Suppl. Figure 3. Consistent with expectations, SMCs, fibromyocytes, and fibroblasts form a distinct cluster, as do macrophages, DCs, and monocytes. Similarly, T cells and NK cells are grouped together, while B cells, plasma cells, and ECs all form their own separate clusters.

To validate the cell type annotations of the atlas, we used two orthogonal methods. First, via label consensus, and secondly with unbiased protein surface markers from CITE-seq data. Three of the datasets (Pauli^35^, Emoto^6^ and Wirka^4^) were annotated independently by an expert and annotations of one dataset were provided by the authors (Jaiswal^36^). The predicted cell types in our atlas were compared to the provided labels. The precision of the annotations is 89.37% in the Pauli dataset and higher than 90% in the three other datasets (see Figure 4 A-D). The confusion matrices also show the mismatch of labels mostly within cell types that are hard to distinguish because of similar transcriptomes, such as NK and T cells or DC, macrophages and monocytes. SMCs, fibroblasts and fibromyocytes are also more difficult to distinguish for the model, as they form one cluster in the UMAP. Overall the consensus is very high with residual uncertainty only between closely related cell types, demonstrating the robustness of the annotations. These results demonstrate a very high consensus between different manual annotations, but it appears impossible to decide which annotation is more accurate in the absence of objective ground truth cell type labels. For a more unbiased validation, we analyzed CITE-seq data^5^ of a sample included in the atlas. The gene expression data was part of the atlas, and the predicted labels were compared to the proteomics surface markers. The available surface markers were grouped by their cell type specificity and visualized across our cell type annotations (see Figure 4 E). The surface markers show highly specific expression patterns across predicted cell types, providing an unbiased validation of our cell type annotation on the protein level.

**Figure 4:**
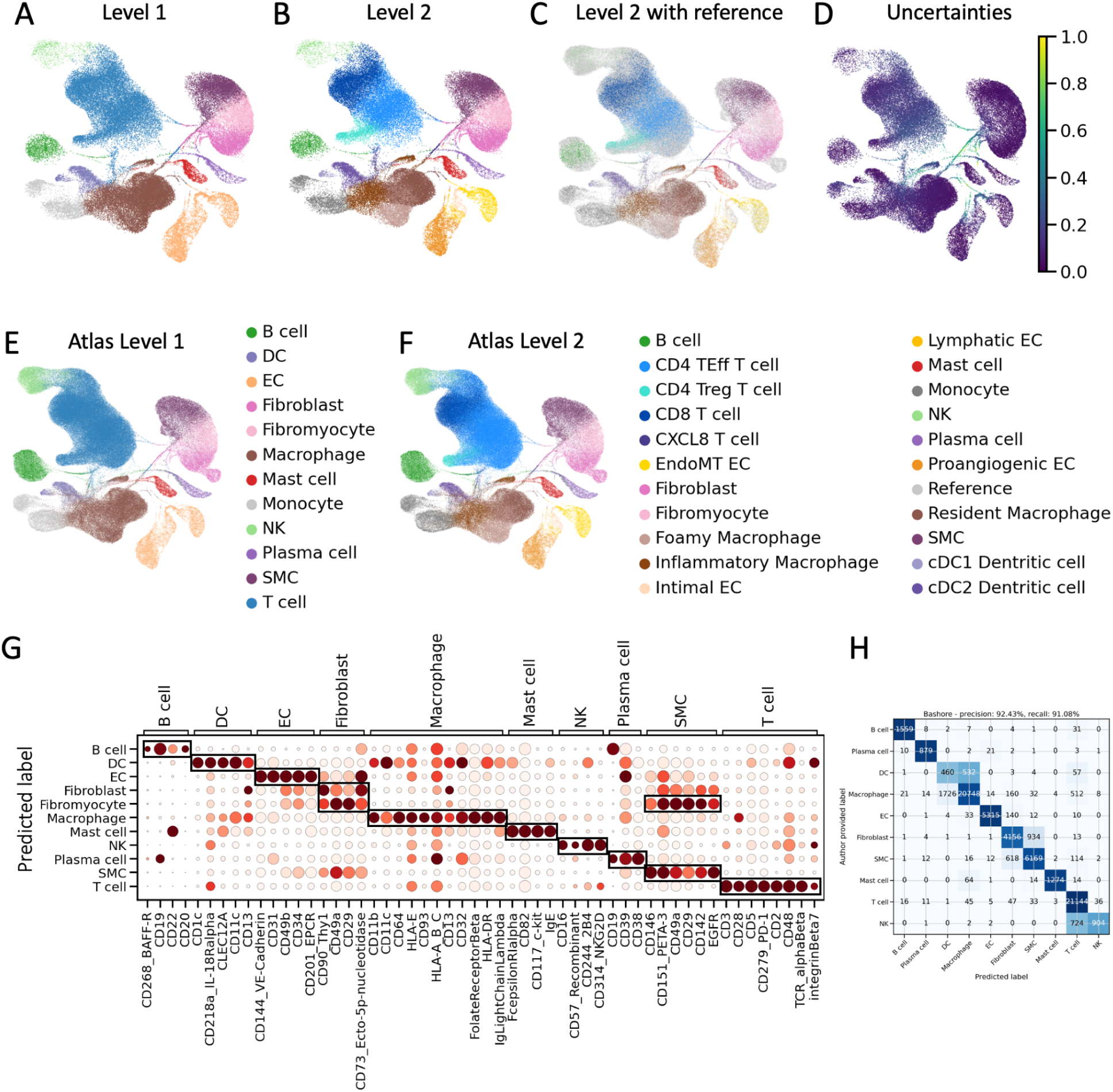
Reference mapping with a dataset consisting of more than 75k cells. Panels A and B show the UMAP projections of the query cells with cells colored by their annotations on level 1 (color legend in panel E) and level 2 (color legend in panel F), while panel C additionally includes cells of the reference in gray. The uncertainties of the predictions are shown by the color code in panel D. Panel G depicts the dot plot of CITE-seq protein expression of surface markers grouped according to cell types (x-axis) for each annotated cell type (y-axis). The highly specific expression pattern of the surface protein is depicted with black boxes. The color coding shows normalized protein expression of the proteins encoded by the marker genes, with light red indicating low expression and dark red indicating high expression. Cells with uncertainty greater than 0.7 are excluded from this analysis. Panel H shows the confusion matrix comparing author provided labels (y-axis) with our predictions (x-axis) with a row normalized colormap to account for differences in cell type abundances.

Overall, this extensive and thorough orthogonal validation demonstrates the robustness of the annotations in the atlas and provides confidence in using it as a reference for future studies.

### Automatic cell type annotation

A key use case of the plaque cell atlas is to automatically annotate and integrate future scRNA-seq plaque datasets by reference mapping. This is key to get robust cell type annotations, which are a prerequisite to compare different data sets to each other. This is necessary for virtually all downstream tasks. To validate the quality of automatic cell type annotations based on our atlas, we analyzed an additional independent carotid plaque dataset^39^ consisting of more than 75k cells, of which more than 25k have additionally been profiled by CITE-seq. We utilized this as a query data set and mapped it to the atlas as a reference using scArches. Figure 4 shows the UMAP projection of the query onto the reference colored by level 1 (Figure 4A) and level 2 cell type annotation (Figure 4B and Figure 4C). In addition to the annotations, the uncertainty of the cell type assignments is provided (see Figure 4D). The annotation shows strong confidence in the majority of cells, while only a small percentage of cells have a high uncertainty. The UMAP projections of the whole atlas including the automatically annotated cells shows that cell types of query cells were assigned consistently with the reference on level 1 and 2 (Figure 4E-F).

To corroborate the accuracy of predicted cell types, we assessed the expression levels of surface proteins measured by CITE-seq. Cells with uncertainty greater than 0.7 were excluded (35 cells out of 24.384). The dot plot indicates highly cell type-specific expression of established surface protein markers^39^ for all cell types (Figure 4G), indicating a high agreement of predictions with the cells’ true identity. Importantly, fibromyocytes express both SMC and fibroblast markers, as these cells are rendered fibroblast-like SMCs^4^.

To demonstrate the consensus of annotations within the field, we compared our predictions with the annotations provided by the authors (see Figure 4H). In a first analysis, the authors’ annotations were harmonized to match our cell types, keeping only the common cell types. Our predictions achieved a precision of 92.43% and a recall of 91.08%, demonstrating high consensus. Similar to the atlas validation, blocks between transcriptionally similar cell types are forming. Additionally, we investigated the annotations of cell types that were not shared between our dataset and the authors’ dataset. In the authors’ dataset, these included neutrophils, whereas in our dataset, monocytes and fibromyocytes were identified. As expected, our predicted fibromyocytes were classified as fibroblasts and SMCs by the authors, while our monocytes were determined as macrophages (Supplementary Figure 4). Neutrophils and cells that were not annotated by the authors were mostly labeled as monocytes.

This validation by unbiased CITE-seq and annotation consensus confirms that the plaque atlas yields highly accurate automatic cell type annotations. To enhance accessibility for researchers annotating their datasets, we provide our atlas as a reference via the user-friendly archmap web server (https://www.archmap.bio/<u>#/</u>) and also offer a solution that does not require data upload, comprising a preconfigured Python script and a Docker container. Both approaches eliminate the need for advanced technical proficiency, enabling researchers of any background to efficiently annotate their datasets.

### Planning future experiments with scPower

The plaque cell atlas also enables to optimize the power of future experiments that aim to compare cell type specific expression between multiple samples of different clinically relevant conditions, such as stable versus unstable plaques. We made use of the scPower framework^14^, which is based on cell type specific gene expression distributions and requires assumptions about the gene expression levels and effect sizes (log fold change) of genes of interest. To enable plaque-specific power analysis, we used the atlas data to learn the parameters of these cell type-specific gene expression prior distributions (Suppl. Figure 5). Moreover, we derived several technical parameters from the Pauli et al.^35^ dataset, which are required as additional input. In the analyses that follow, we specified a fixed budget as well as the costs of typical sequencing runs (see Methods).

To evaluate the power to detect differentially expressed gene signatures, we formulated assumptions for three scenarios with different average effect sizes of differential gene expression: a pessimistic, neutral, and optimistic scenario with fold changes (FC) 1.1, 1.5 and 2.0 respectively. We evaluate the average power for a gene signature composed of 20 genes, which are expressed among the top 100, 1000 and 5000 genes, respectively. For abundant cell types the power is generally high, while lower abundance cell types show almost no power (Figure 5). As expected, higher effect size leads to an increase in power. Also, the power increases if the signature is composed of more highly expressed genes corresponding to smaller expression ranks. Notably, the most disease-relevant cell types according to the current literature such as T cells, macrophages and SMCs^4,5^ are abundant enough to yield a high power. In case an investigation of rare cell types such as DCs, B cells or mast cells is desired, we recommend sorting with FACS prior to sequencing to enrich these cells enough to have sufficient power to detect differentially expressed genes.

**Figure 5:**
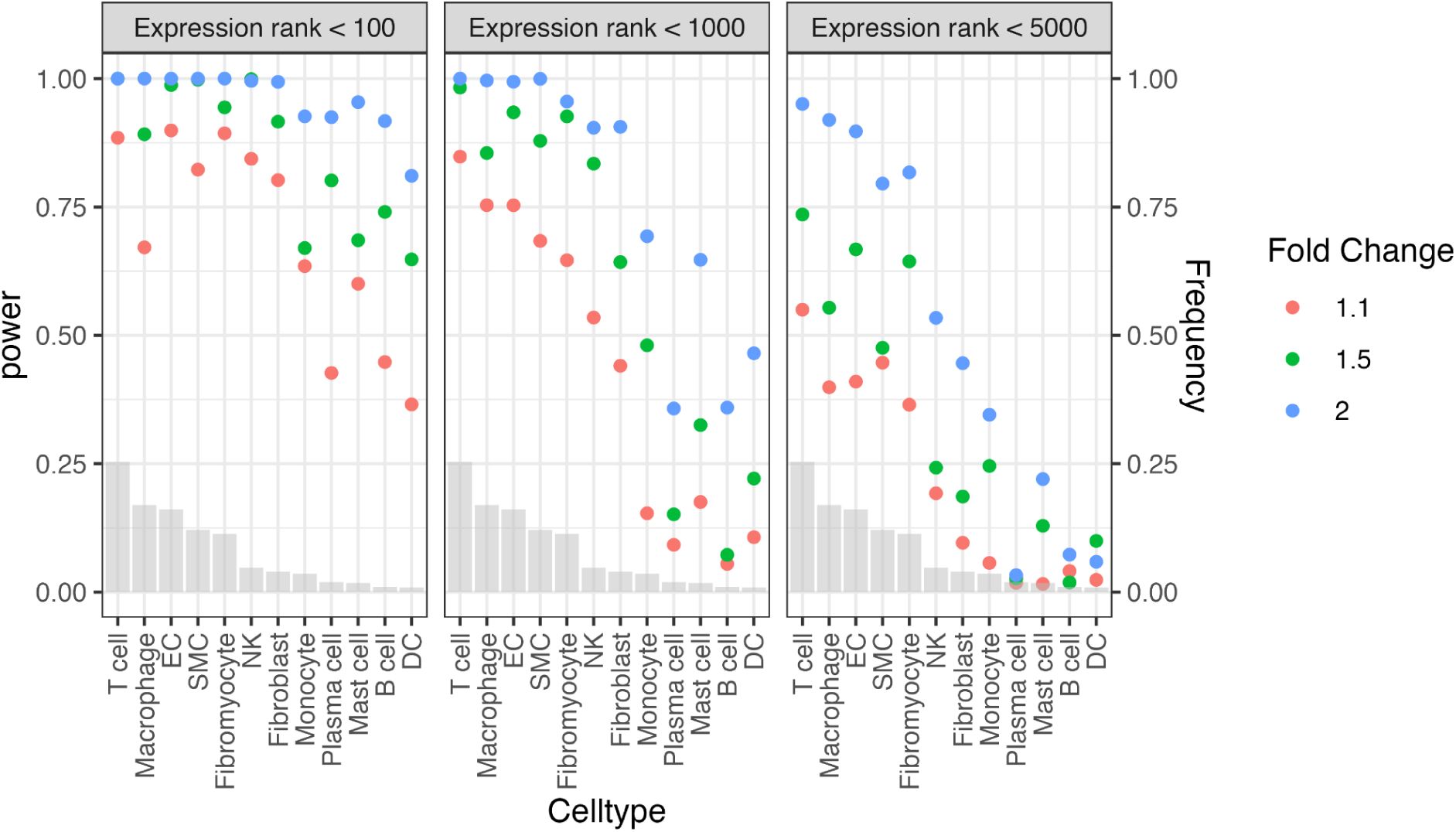
Power analysis utilizing the scPower framework. The plot shows the statistical power of gene signatures made up of 20 genes sampled from the top 100, 1000 and 5000 ranking genes in expression with a pessimistic (FC 1.1), neutral (FC 1.5) and an optimistic (FC 2.0) scenario per cell type. The gray bars depict the expected frequency of the cell types in the scRNA-seq study.

This framework provides a valuable resource for the research community to plan future experiments more efficiently and cost-effectively. A web-based dashboard (https://scpower.helmholtz-muenchen.de/) allows users to specify their own parameters, e.g. expected cell type abundances, samples, cells or financial budget and others. Additionally, it is possible to obtain power calculations for very specific gene sets or pathways that can be provided based on pilot experiments. This enables researchers with less technical proficiency to first use the web-based scPower tool to plan their experiments and subsequently automatically annotate their data using the web-based archmap tool described above. This user-friendly web-based workflow makes the field of scRNA-seq more accessible to the cardiovascular research community.

### Mapping on diverse tissues

Fibromyocytes and macrophages are frequently discussed as relevant cell types for plaque progression^4,66^. Having defined these plaque-specific cell types in our atlas, we next wondered if these cell types are exclusively found in atherosclerotic plaques. Therefore, we investigated 455,953 cells of a total of 23 different scRNA-seq datasets from various organs^67^ and assessed their similarity to the cells in the atlas using the automatic mapping tool. Four exemplary organs are showcased in Figure 6. Interestingly, fibromyocytes were only found in the vasculature in significant numbers (n > 20), highlighting that cells that were labeled as fibromyocyte are distinct enough to not be detected in other tissues. Of note, the vascular cells in the Tabula sapiens data set were sampled from individuals with CAD. To investigate the presence of fibromyocytes in healthy vascular tissues, we mapped a dataset of healthy arteries^22^ onto our atlas (see Suppl. Figure 6). The analysis revealed a significant abundance of fibromyocytes in these healthy vessels, indicating that these cells are not exclusive to plaques but are a normal component of the vasculature, in line with the results of previous studies^23^.

**Figure 6:**
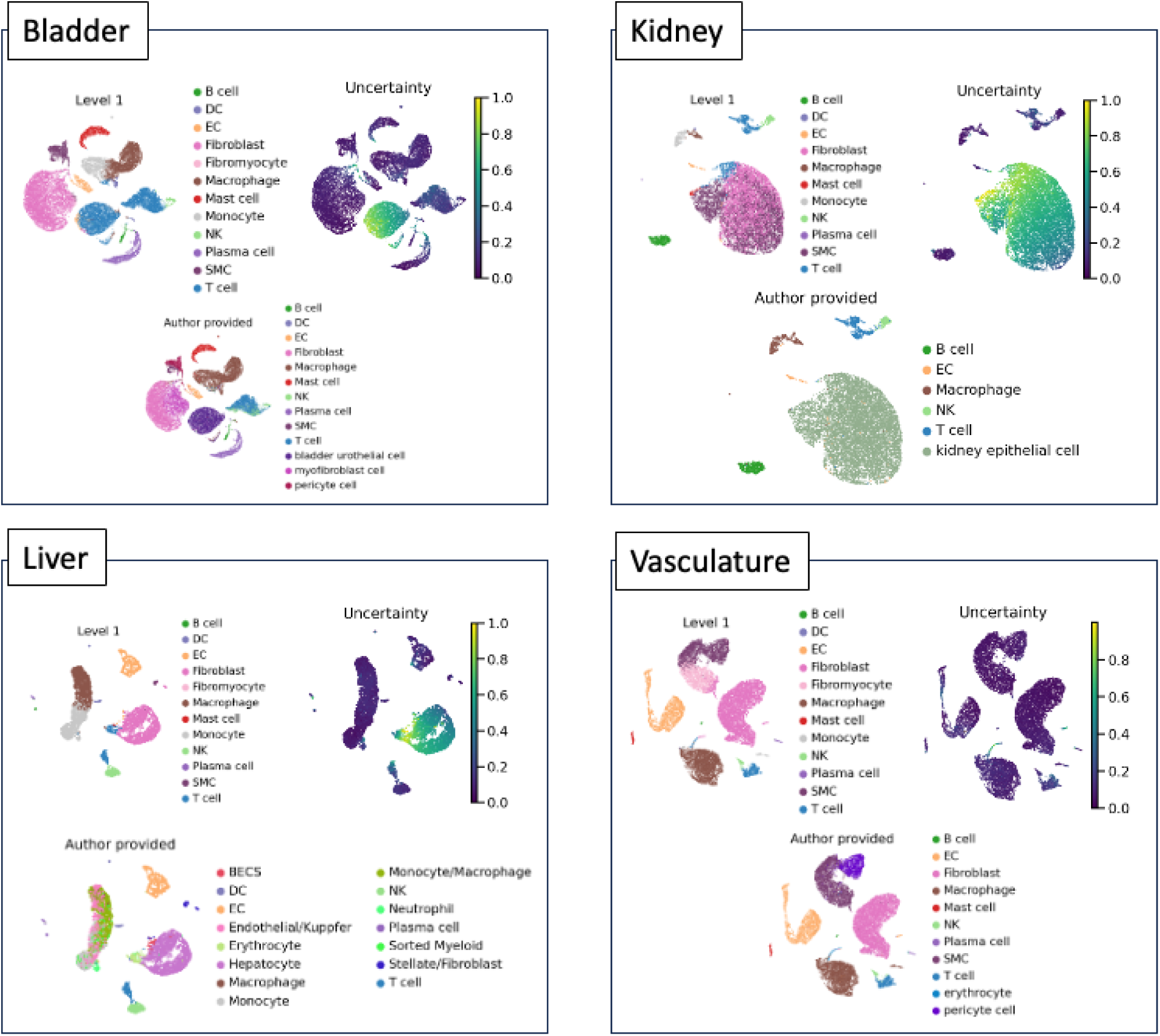
Mapping Tabula Sapiens organs scRNA-seq datasets on plaque atlas. Four exemplary mappings of the Tabula Sapiens dataset on the plaque atlas. For each organ, the predicted level 1 cell type is shown on the top left panel, the models uncertainty on the top right and the free annotations provided by the Tabula Sapiens authors on the bottom.

We also screened for foamy macrophages in other tissues and found them highly abundant in accordance with the literature in the liver^68^ and lung^69^ but also in the vasculature (see Suppl. Figure 7). In the original analyses these cells were mostly annotated as macrophages and their foamy phenotype was not recognized.

As expected, the uncertainty for annotating cells originating from vascular tissues is the lowest, because the atlas includes the surrounding tissues of the plaques as well. In all other organs we observed clusters of cells with very high uncertainty. Further analysis revealed that these clusters are organ specific cells, such as bladder urothelial cells in the bladder, kidney epithelial cells in kidney, and hepatocytes in liver. In the vasculature dataset there was a small cluster of uncertain cells, which were identified as erythrocytes that are not part of our atlas. Simultaneously, the clusters with low uncertainty were correctly assigned by the automatic mapping to the cell type annotated by the authors (see Suppl. Figure 8). Together this highlights the robustness of our atlas and indicates that atherosclerosis associated cell types can also occur in healthy subjects.

### Deconvolution of bulk RNA samples

Another downstream task that can be achieved with the plaque cell atlas is the deconvolution of bulk RNA samples. Because of the complexity and costs of scRNA-seq experiments, sample sizes of studies are usually limited, while the information per sample is very abundant. Because bulk RNA experiments are cheaper per sample, cohort sizes are bigger, the number of identified transcripts is much higher, while information of cellular origin is obviously lacking. Deconvolution approaches combine both worlds, by using the abundant cell type specific gene expression information from scRNA-seq as priors, to deconvolute bulk samples into a mixture of cell type specific expression profiles and their respective cell type abundances. We used BayesPrism^17^ to deconvolute a dataset of 236 (202 after QC, see methods) bulk RNA seq samples from carotid artery plaque tissue^35^. For compositional data analysis the estimated abundances were centered and log ratio transformed (CLR).

The carotid plaque samples are classified into early or advanced (late) lesions of the carotid artery (see methods). This enables us to investigate differences in cell type composition between early lesions and diseased carotid arteries. Structural cells such as fibromyocytes, SMCs, ECs, and fibroblasts together with macrophages were most abundant (see Fig 7A). Stratifying by plaque status reveals significantly more SMCs (t-test on CLR values: P=6.1e-09) in early lesions than in advanced lesions and vice versa significantly more fibroblasts (t-test on CLR values: P=1.8e-08) in late, progressed lesions. This indicates the high cellular plasticity of SMCs and their transition from SMCs to fibroblast-like cells^4^. Interestingly, there appears no significant difference in fibromyocytes between early and advanced lesions in carotid lesions. It can be speculated that not a change of fibromyocyte abundance, but rather a shift of their gene expression is associated with plaque progression. This is in line with the observation that fibromyocytes can be detected in artery samples without late lesions in the Tabula Sapiens data set and healthy arteries in the Hu et al. dataset^22^. Additionally, significantly more macrophages (t-test on CLR values: P=5.7e-05) are observed in advanced lesions validating the infiltration and increased activity of macrophages in more progressed atherosclerotic lesions and the immune system’s contribution to disease acceleration^8,70^. Finally, a small but significant (t-test on CLR values: P=4.3e-02) difference in EC abundance is detectable.

**Figure 7:**
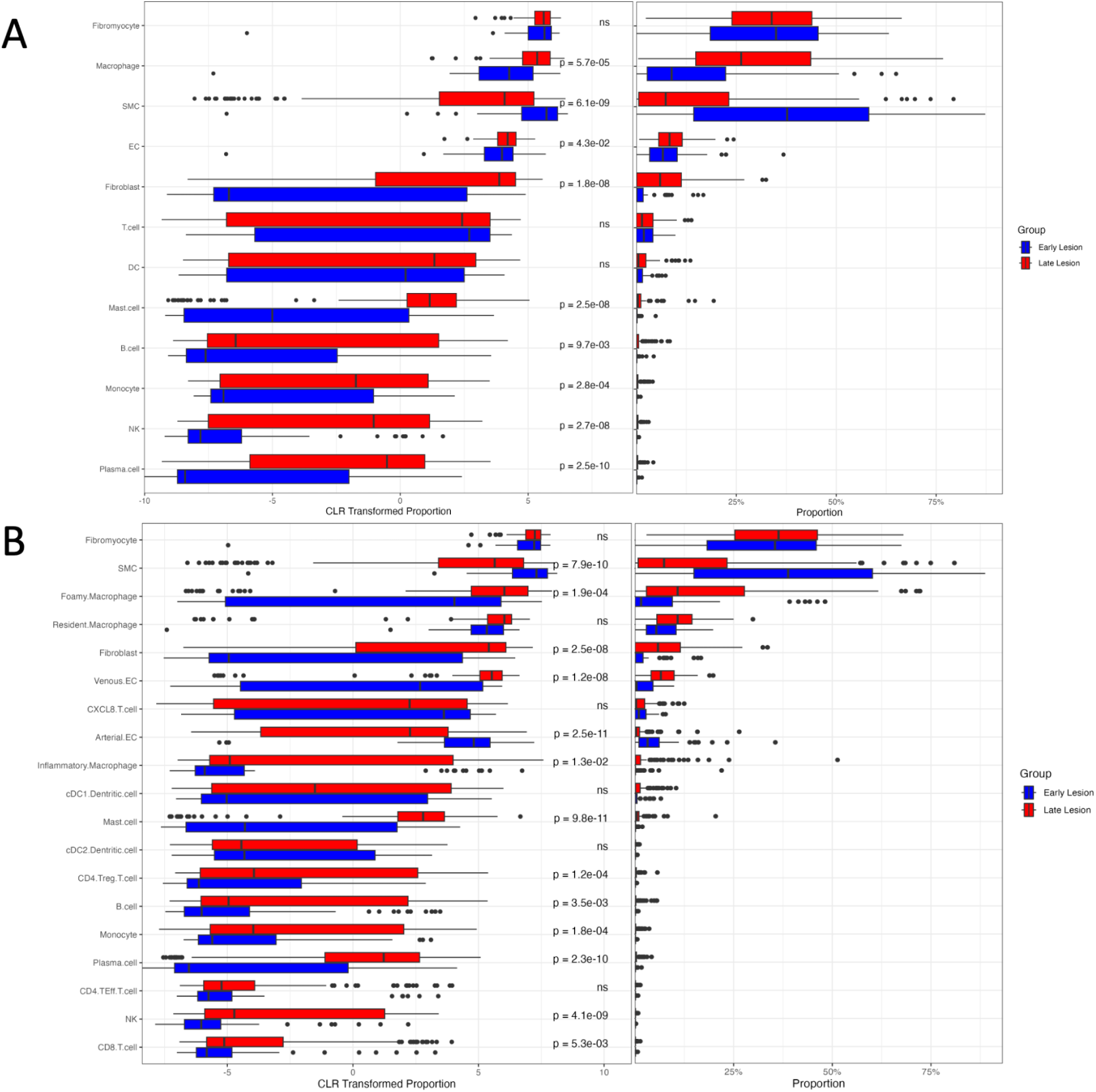
Deconvolution of bulk RNA-seq samples using BayesPrism and the plaque atlas. The cell type proportions (right panels) were centered and log transformed (CLR) to show differences in small abundances better (left panels). A shows the stratified deconvolution results using the level 1 cell types in the atlas as the reference. B depicts the deconvoluted abundances of the bulk samples using the finer grained level 2 cell types in our atlas. ECs were re-annotated to venous and arterial ECs.

To highlight these differences for finer granular cell types, we selected all EC cells of the atlas using our level 1 annotations. Subsequently, we individually corrected them for batch effects on the counts, and clustered the ECs for arterial and venous origin based on varying marker gene expression. While the vast majority of proangiogenic ECs were mapped to venous ECs and most EndoMTs mapped to arterial ECs, the intimal and lymphatic subtypes mapped roughly equal to arterial and venous ECs (see Suppl. Fig 9). The possibility to easily reannotate level 1 cell types of the atlas with finer grained cell types of interest, depending on the downstream task, demonstrates the flexibility of the atlas. For the subsequent deconvolution, we used the level 2 cell types with finer granular macrophage subtypes such as foamy, resident and inflammatory macrophages (see Fig 7B). The aforementioned differences in SMCs and fibroblasts are still visible while new details are revealed. The newly annotated EC subtypes show highly significant differences between early and advanced lesion tissues. While the venous ECs are more abundant (t-test on CLR values: P=1.2e-08) in more advanced plaques, arterial ECs present the opposing abundance pattern (t-test on CLR values: P=2.5e-11) in early lesion tissues compared to advanced, late lesions. This interesting observation confirms the notion that these EC-rich neovessels in advanced plaques are likely the entry point for immune cells that infiltrate the tissue and contribute to lesion progression and instability. This becomes particularly relevant when EC barrier function becomes impaired, neovessels start to leak, and intraplaque hemorrhages occur. The stratified macrophage subtype abundances show results that are well in line with previous literature, as there appears no significant difference in resident macrophage abundance in advanced lesions, validating their likely passive role in atherosclerotic progression. On the other hand, inflammatory and foamy macrophages are substantially enriched (P-value) in more advanced lesions, reflecting the important role of macrophage infiltration and activation in late-stage atherosclerotic plaques, as described in previous studies^8,70^.

Overall, bulk to single cell RNA-seq deconvolution demonstrates the utility of a robustly annotated atlas by highlighting disease-relevant biological processes and comprehensively dissecting cell subtype abundance and relevance for atherosclerotic plaque formation and progression.

## Discussion

The plaque atlas showcased in this study represents the most comprehensive curation of single-cell RNA sequencing (scRNA-seq) datasets of atherosclerotic lesions to date^23,24^. This atlas, extensively validated through annotation consensus, protein measurements, and the illustration of known biological processes, serves as a robust reference for future studies. It provides a foundation for integrating new research questions and advancing our understanding of atherosclerosis.

Previous integration efforts for single cell data from plaque tissue ^23,24^ were often constrained by limited scope, validation, and usability. In contrast, our atlas encompasses all major lesion locations, offers thorough orthogonal validation, and supports user-friendly interactions through an interactive web interface. Despite having defined fine-grained level 2 cell type annotations, we recognize that the consensus on these annotations can vary depending on the specific research question. Often, these specific subtypes are transcriptionally highly similar, which makes it difficult to unambiguously distinguish them, especially for smaller level 2 cell type populations (e.g., CXLC8 positive T cells). Hence, we recommend using level 1 annotations initially, followed by sub clustering into level 2 cell types of interest, as demonstrated in our bulk deconvolution workflow, which yielded results in line with previously published studies from the cancer field^17^.

The scPoli model, integral to this atlas, is trained on predefined cell types, limiting its predictive capacity to those included in the study. To mitigate this limitation, we included the most commonly studied cell types, while excluding those with less clear distinction or abundance. Notably, the model provides uncertainty metrics for newly mapped cells, suggesting that clusters of cells with high uncertainty in new datasets could indicate the presence of yet unidentified cell types not included in the atlas, as demonstrated with the non-plaque tissue mappings.

Overall, this atlas not only serves as a robust foundation for future research but also enhances accessibility for newcomers to the field, by making it easy to use. Moreover, it can be integrated into the training of foundation models^19–21^, thereby expanding the dataset corpus to include plaque tissues. This inclusion is crucial, as current models often underperform in out-of-distribution tasks, reflecting an overly homogeneous training corpus.

We demonstrated three primary downstream tasks: automated cell type annotation and cross-organ analysis, power analysis and study planning, and bulk RNA sample deconvolution. For future studies, the atlas can be used in additional downstream applications, such as integrating scRNA-seq data with spatial transcriptomics datasets and conducting cell type-specific genome-wide association studies using prior atlas information^23^. Given the availability of specially designed bulk RNA-seq experiments, the atlas can facilitate the deconvolution of phenotype-specific signals or survival analysis. Reference mapping, including healthy reference atlases, can highlight differences between healthy and diseased tissue samples at the single-cell level, as recently emphasized^71^. This underscores the necessity of an atlas comprising exclusively diseased samples.

In summary, this comprehensive and robust atlas offers numerous possibilities for downstream applications and provides an invaluable resource for uncovering novel cellular processes in atherosclerosis.

## Data/code availability

The Python and R code to reproduce the results is available at https://github.com/kotr98/reproducibility-plaque-atlas. The python script and docker container for the automatic cell type annotation is available at https://github.com/kotr98/plaque-atlas-mapping and https://github.com/matmu/cell_type_annotation.

The data used in this study is all publicly available under the provided reference. The core atlas (without the Bashore et al. mapping) is uploaded to the CELLxGENE portal at https://cellxgene.cziscience.com/collections/db70986c-7d91-49fe-a399-a4730be394ac.

## Methods

### Preprocessing of data sets

In all datasets QC was applied on the uncorrected counts. Doublets were tagged with *scDblFinder*^26^ with the sample as the batch parameter and the counts are corrected for ambient RNA with *celda*^72^ using the sample as the batch parameter and rounded to integers. The uncorrected counts and celda corrected counts were kept as separate layers.

#### Pan et al.^37^

The three samples were downloaded from GEO and came already filtered with 200 < #genes < 4000, maximum counts of 20.000 per cell and mitochondrial gene count percentage lower than 10%. The three samples were outer joined with the *concat* method of the anndata^73^ package.

#### Alsaigh et al.^38^

The dataset was downloaded from GEO and only plaque samples (barcode suffixes 2, 4 and 6) were taken and adjacent tissue samples excluded. The samples were renamed from 2, 4 and 6 to 1, 2 and 3 respectively. Cells were filtered on the uncorrected counts according to the original publication with 200 < #genes < 4000 and percentage_counts_mitochondrial > 10%. Additionally, we filtered out genes expressed in less than 3 cells.

#### Fernandez et al.^5^

The dataset was downloaded from GEO and each sample is read in with *scanpy*^74^ and outer joined with the *concat* method from anndata. As sample 6 is a CITE-seq sample it also includes surface protein markers. These were removed from the dataset as the atlas is only based on gene expression. We applied our own filtering on the uncorrected counts and filtered out cells with less than 200 expressed genes, 500 < #counts < 40000 and more than 10% mitochondrial gene percentage. Genes were filtered out that are expressed in less than 3 cells.

#### Pauli et al.^35^

The dataset was preprocessed with Seurat with nFeature_RNA > 200 & nFeature_RNA < 10000 & nCount_RNA > 1000 & nCount_RNA < 50000 & percent.mt < 20. Only the diseased samples were taken and the adjacent samples excluded.

#### Dib et al.^8^

The dataset was downloaded from GEO and the sample ids extracted from the barcode suffixes. The sample ids were changed from 5 to 4, sample 6 to 5 and sample 7 to 6, but kept sample id 1, 2 and 3 the same. Cells were filtered on the uncorrected counts with more than 400 genes expressed, 1000 < #counts < 30000, and pct_counts_mt < 10%. Genes that are expressed in less than 3 cells were filtered out.

#### Slysz et al.^9^

The samples were downloaded from GEO. Femoral and carotid samples were loaded in and concatenated with *concat* of anndata into a femoral and carotid anndata object. For both objects the cells were filtered on the uncorrected counts according to the authors with 200 < #genes < 10000, 200 < #counts < 10000 and pct_counts_mt < 10.

#### Jaiswal et al.^36^

The dataset was downloaded from GEO. We selected the “Fresh_ROB_2026” and “Fresh_DTAN_4047” samples and concatenate them into one dataset with *concat* from the anndata package. Metadata was also available and cells which had “Removed by QC” as cell types were removed. To be as consistent as possible we converted the provided ensembl gene ids in this dataset into gene names using the mapping created out of the Fernandez et al. dataset^5^. 4250 genes found no mapping and were removed and kept 32.251 genes. Cells were additionally filtered on the uncorrected counts by us with min_genes = 200, 500 < #counts < 40.000 and pct_counts_mt < 10, while genes that are expressed in less than 3 cells were excluded as well.

#### Emoto et al.^6^

The dataset was downloaded from GEO and the SAP and ACS samples were preprocessed according to the author with min_genes=500, min_cells=3, pct_counts_mt < 8 and max_genes = 5000. Subsequently, both datasets were concatenated with *concat* from the anndata package. The ACS samples are termed sample 1, while the SAP samples are termed sample 2.

#### Chowdhury et al.^7^

The dataset was downloaded from GEO and came already filtered with the following criteria: Genes were filtered that are expressed in less than 5 cells. Cells were filtered with more than 38% pct_counts_mt. Cells for 10x v2 samples were filtered with min_genes = 300 and max_counts = 15.000, while 10x v3 samples were filtered with min_genes = 500 and max_counts = 20.000. The blood samples were excluded and only the plaque samples were kept. The sample ids were changed from alphabetical to numerical according to the letters position in the alphabet.

#### Wirka et al.^4^

The dataset was downloaded from GEO and came already filtered with the following criteria: Genes expressed in less than 5 cells were filtered out. Cells were filtered with pct_counts_mt < 7.5% and kept with 500 < #genes < 3500. There were duplicate barcodes which we investigated more closely. They had different gene expressions, hence we assumed they are different cells and made the barcodes unique.

#### Bashore et al.^39^

The dataset was downloaded from GEO and only the gene expression data from the samples was read in and concatenated with *concat* of the anndata package. It was filtered according to the authors with 200 < #min_genes < 6000, max_counts = 40.000, min_cells = 3 and pct_counts_mt < 30. For this dataset the doublet tagging ambientRNA correction was applied shortly before the reference mapping.

### Manual annotation of samples

Because samples within datasets can also entail batch effects due to the data collection or other factors, we annotated and integrated on a sample level. We manually annotated sample 1,2 and 3 of Alsaigh et al., sample 2 and 3 of Pan et al., sample 6 and 8 of the Slysz et al. femoral dataset and sample 5,6,7 and 8 of the Wirka et al. dataset. For all datasets we performed *scran*^75^ normalization with an initial clustering of total counts normalization with 1e6 as target sum, log1p normalization, PCA, neighborhood calculation on 30 PCs and Leiden clustering with a resolution of 0.22. The resulting size factors were used to normalize the counts. Subsequently log1p transformation is applied, top 2000 highly variable genes selected, PCA and neighborhood calculation on 30 PCs. Then UMAPs are calculated for each sample and clusters were manually annotated with the level 1 marker genes. Doublet clusters were annotated as doublets. Clusters with no distinct marker gene patterns were annotated as unknown. In cases of uncertainties of cluster annotations, these clusters were sub clustered and annotated on a finer resolution.

### Benchmark of integration methods

All annotated samples are concatenated into one dataset and the gene names are mapped to Ensembl ids to solve the problem with changing gene names and aliases. Duplicate genes after the mapping were aggregated. The resulting dataset was normalized with *scran* using the same parameters as in the manual annotations and log1p transformed. The sample ids were suffixed with the dataset name to make the sample ids unique. Unknown and doublet cells were removed. It was preprocessed with the *reduce_data* method with sample id as batch_key of the *scib*^27^ package. Highly variable genes were selected and cells with zero counts were removed. Additionally, cells with the duplicate gene expressions were removed as well. The methods that are benchmarked are scVI^28^, Harmony^29^, LIGER^30^, scANVI^31^, scGen^32^, scPoli^33^ and baseline PCA. For scGen, the *scib* implementation was used on the log-normalized counts with the sample id as the batch key and the manual annotations as cell types. For *scVI* the default parameters and counts are used. Subsequently the scVI model is fine-tuned with *scANVI* with the cell type labels. For *scPoli*, the default parameters are used, but the loss is changed to ‘mse’ and calculated on the log-normalized counts. The *Harmony* is calculated on the PCA embedding with the *harmony-pytorch* package. *LIGER* embeddings are calculated following the tutorial on *scib-metrics*^27^. The resulting embeddings are benchmarked with the *scib-metrics* benchmarker using the sample id as batch key and cell types as label key.

### Atlas integration

To generate the whole atlas all preprocessed datasets except Bashore et al. were concatenated with *concat* from anndata. The sample ids were made unique by suffixing the dataset. Subsequently the gene names were harmonized by mapping them to ensembl genes and aggregating duplicates. Duplicate and zero count cells were removed and the whole dataset was *scran* normalized and log1p transformed with the parameters described earlier. Manual annotations were transferred from the benchmark subset. For the preprocessing of the whole object, the same pipeline as in the benchmark was applied. A split of reference and query dataset was performed where the manual annotations serve as the reference. A scPoli model was trained on the reference dataset log-normalized counts with the sample as the condition key and the cell type annotations as cell type key. An embedding dimension of 10 was chosen and trained with a mse reconstruction loss. For everything else default parameters were used. Subsequently, the query dataset was mapped to the reference using the pretrained reference scPoli model and did transfer learning with scArches and new labels including uncertainties predicted.

### Level 2 annotations

For finer grained cell types we used the level 1 predictions from the atlas integration. We selected all cells of a certain cell type, took the rounded corrected counts layer and applied our standard preprocessing pipeline to log-normalize it. Afterwards, we used *Harmony* without cell type annotations to integrate the samples to eliminate potential batch effects and conserve as much biological signal as possible. The resulting dataset was clustered with the Leiden algorithm and manually assigned to cell types with our level 2 marker genes. This was done for macrophages, dendritic cells, T cells, ECs and NK cells. We were also looking for potential mis-classifications and corrected them by assigning the correct cell type. Subsequently, the scPoli model was trained again with the same procedure as in the level 1 training, but with the newly assigned level 2 annotations. We also added the level 1 cell types again by inferring a new column in the dataset using the level 2 cell types.

### Validation of annotations (expert and CITE-seq)

To validate the annotations, an expert annotated three datasets (Pauli^35^, Emoto^6^ and Wirka^4^) independently from us without our help or interaction using his own methods and judgment. One dataset^36^ was annotated by the authors of the original publication. To compare the cell type annotations with our predictions, they needed to be harmonized. For this, we mapped the level 2 cell type predictions to level 1 cell types and did the same to the provided annotations from the expert and authors. The intersection of cell types between our predictions and the author/expert provided annotations were used to calculate confusion matrices and precision/recall. The precision and recall was calculated per cell type and then weighted according to their abundance to yield an overall precision and recall per dataset. For the CITE-seq dataset we looked at the protein measurements and used the *dotplot* function in *scanpy* to plot our predicted cell types in these samples to the surface markers that we grouped into cell types, while scaling the expression between 0 and 1 within each surface protein (default).

### Automatic cell type annotation on Bashore dataset

The gene expression data of the Bashore et al.^39^ dataset was used to validate the automatic cell type annotations. We made the sample names unique, mapped the gene ids to ensembl ids with our mapping, removed 22 non-mapped genes and aggregated duplicated genes (same workflow as before). Our standard *scran*-log1p normalization/transformation was applied. The dataset was then subsetted for the 2000 genes used in the scPoli model, where 9 genes were not in the query dataset. The missing information was filled with zeroes. This resulted in 77.112 cells that were loaded into the reference model and trained with scArches. Finally, the level 1 cell types were inferred from the level 2 prediction and cells with uncertainty higher than 0.7 removed. The same pipeline for validation with confusion matrices and protein surface markers as in the previous section was applied, where the author provided us with the cell type labels. Cells that were not given a label by the authors were labeled as “unknown” by us.

### Power analysis with scPower

To calculate the gene expression priors the uncorrected counts and level 1 cell type annotations were used. The gamma and dispersion fits were calculated per cell type. The matrices were subsampled using multinomial sampling to simulate 25%, 50% and 75% total counts. Because the T cell matrix was too big in memory, we used a more memory efficient subsampling method that uses sparse matrices instead of dense matrices. Subsequently, the gammas and dispersion priors were fitted using the tutorial in the vignette of *scPower*. To model the transcriptome mapped reads to UMI relationship we used the Pauli et al. dataset as priors where we had CellRanger outputs. The same holds for the cell type frequency priors, for which we used the predictions of the atlas in this dataset. We assumed 22 samples with 200 cells each, a read depth of 334.015 and mapping efficiency of 43% which were the mean parameters in our samples. All of these parameters can be easily changed in our interactive web-based dashboard according to your experimental setup.

### Sample collection bulk RNAseq data

Sample collection and preparation for sequencing was performed as described before^35^. The classification of human carotid artery samples into early and late-stage atherosclerosis was initially performed visually during the cutting and preparation process. This means the identification and separation of advanced plaque parts with small lumen versus large lumen without visually detectable plaque. The classifications were then confirmed through subsequent histomorphological analysis of FFPE-processed adjacent sections. Lesions were categorized according to AHA guidelines^76–79^ as previously described^35^. Samples classified as I, II and III were designated as early lesions, while those classified as V to VIII were assigned late lesions.

### Preprocessing bulk RNAseq data

We performed adaptor clipping and quality trimming of sequences using Fastp v0.23.2 (https://github.com/OpenGene/fastp). Subsequently, reads were aligned to the GENCODE v40 GRCh38 reference transcriptome using the transcript quantifier Salmon v1.6.0 (https://salmon.readthedocs.io/). The R package tximeta was employed to combine Salmon transcript quantifications with sample data and to summarize transcript quantifications at the gene level.

To assess data quality at both the read and alignment levels, we utilized FastQC v0.11.9 (www.bioinformatics.babraham.ac.uk/projects/fastqc/) and MultiQC v1.13a (https://multiqc.info/) to generate statistics before and after trimming. Outlier samples were detected by performing a principal component analysis (PCA) on variance-stabilizing transformed counts for the 500 genes with the highest variance, using the R package DESeq2.

Based on the quality control metrics, we removed samples that met the following criteria: PCA outliers (determined by visual inspection), percentage of mapped reads < 75%, percentage of duplication > 60%, and number of mapped reads < 10,000,000. After filtering, a total of 202 samples remained.

### Deconvolution of bulk RNA-seq experiments with BayesPrism

To deconvolute the bulk samples we used BayesPrism and closely followed their tutorial provided in the GitHub repository (https://github.com/Danko-Lab/BayesPrism/). The uncorrected counts of the atlas including the cell type annotations were loaded and Ensembl IDs were used in both the scRNAseq reference and the bulk matrix. Genes were first filtered with the cleanup.genes function with default parameters, then filtered for protein coding genes with select.gene.type and finally signature genes were calculated with get.exp.stat and subsequently filtered with select.marker. For the level 1 deconvolution the level 1 cell types were used as the cell.type.labels and the level 2 cell types used as the cell.state.labels. For the level 2 deconvolution both the cell.type.labels and the cell.state.labels were set to the level 2 cell types in the atlas. Finally, the theta values for each cell type per sample are used for the figures. The cell types are sorted according to their mean proportions across all samples. The abundances were centered and log transformed with the clr function of the *compositions*^80^ package. For significance testing a t-test with default parameters was applied on the CLR transformed values.

## Acknowledgements

K.T. is supported by the Helmholtz Association under the joint research school “Munich School for Data Science – MUDS”. L.M. is supported by the ERC Consolidator Grant LongTX (under the grant agreement number 101088370), the Bavarian State Ministry of Health and Care through the DigiMed Bayern project on P4 medicine, and the DFG-funded TRRs/CRCs 267 (Non-coding RNAs in the cardiovascular system) and 1123 (Novel targets in atherosclerosis). M.H. is supported by the Chan Zuckerberg Foundation (2019-202666, 2021-237882) and the DZHK (German center for cardiovascular research) projects 81Z0600106 and 81Z0600105.

