## Supplementary Figures for "Integrated single-cell atlas of human atherosclerotic plaques"

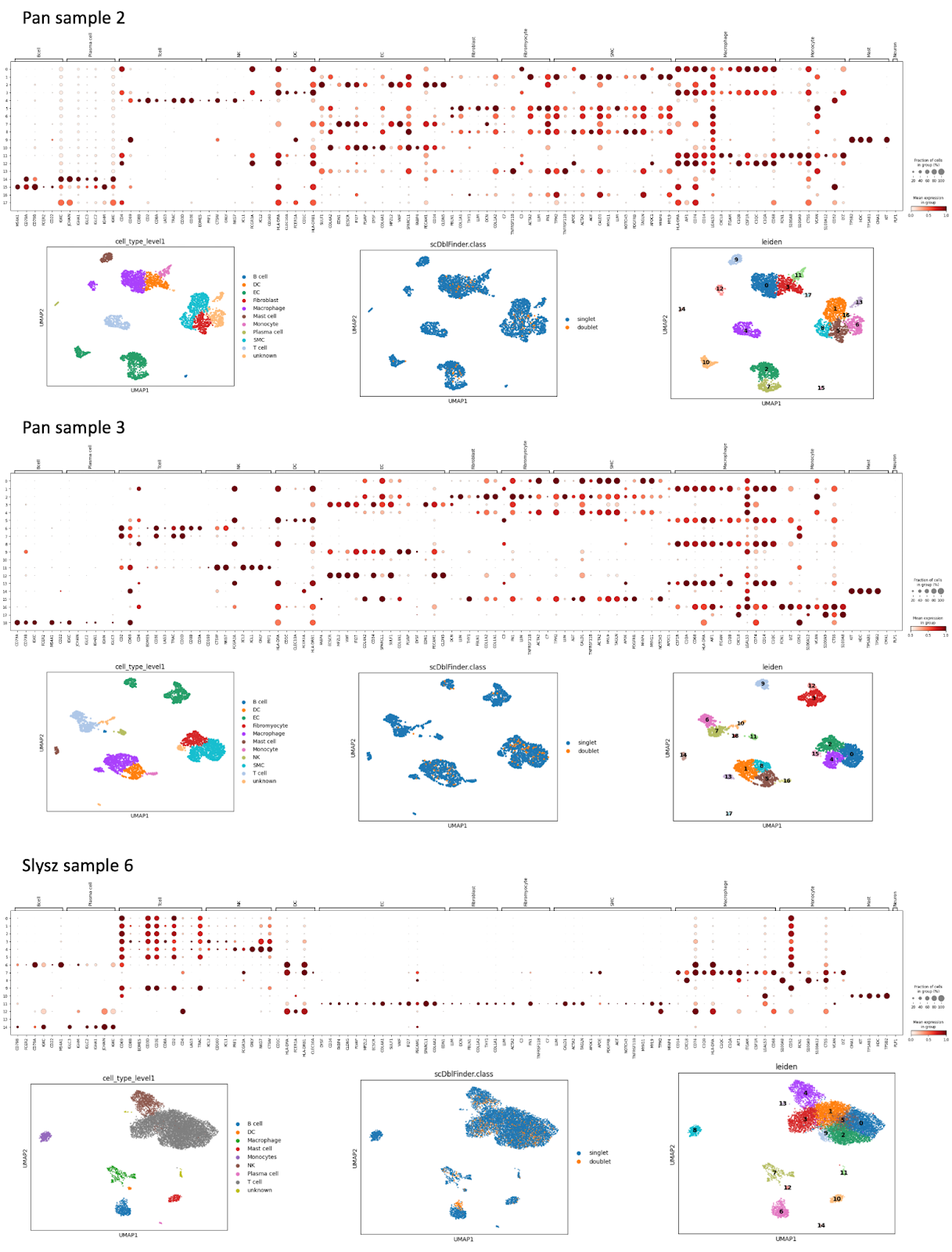

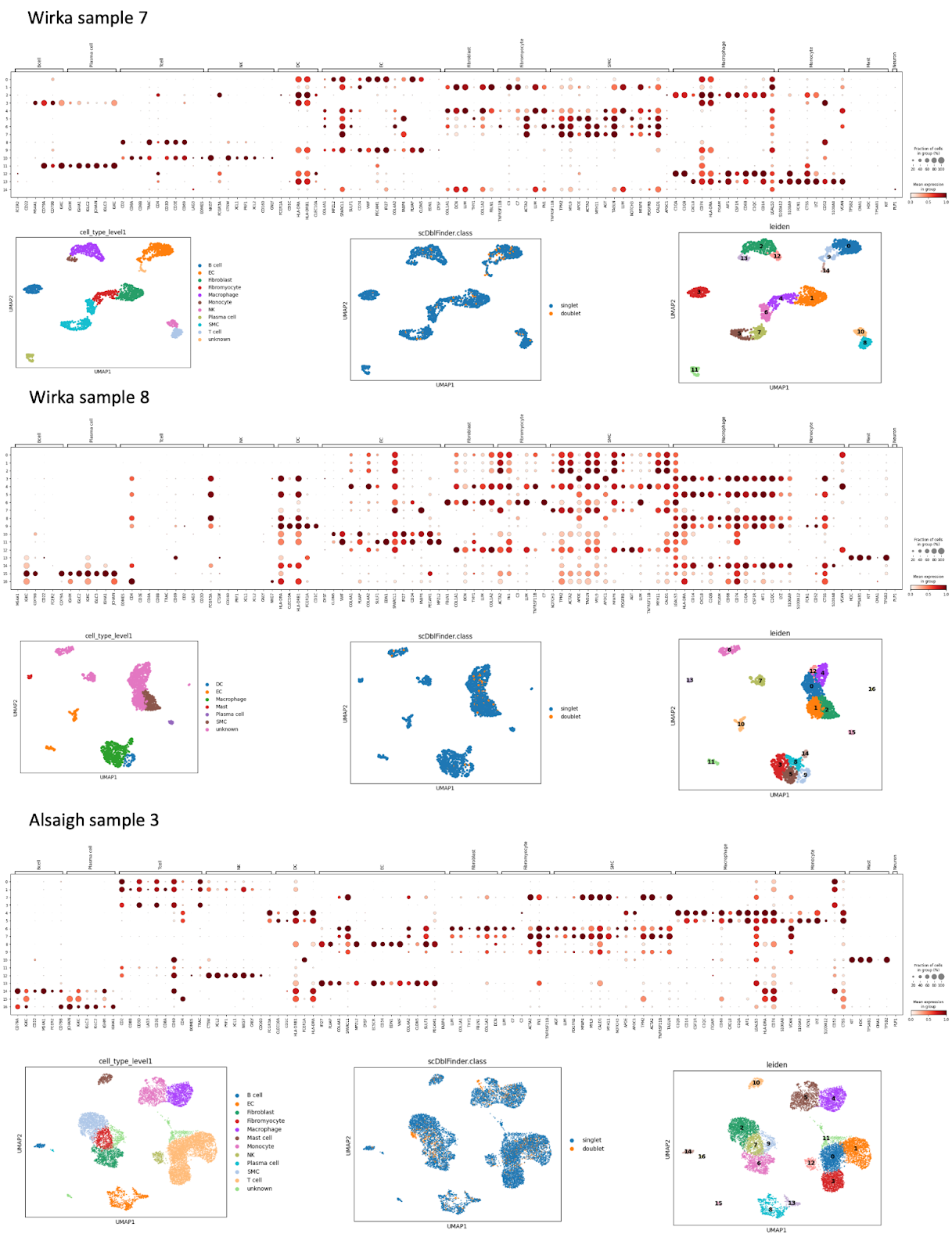

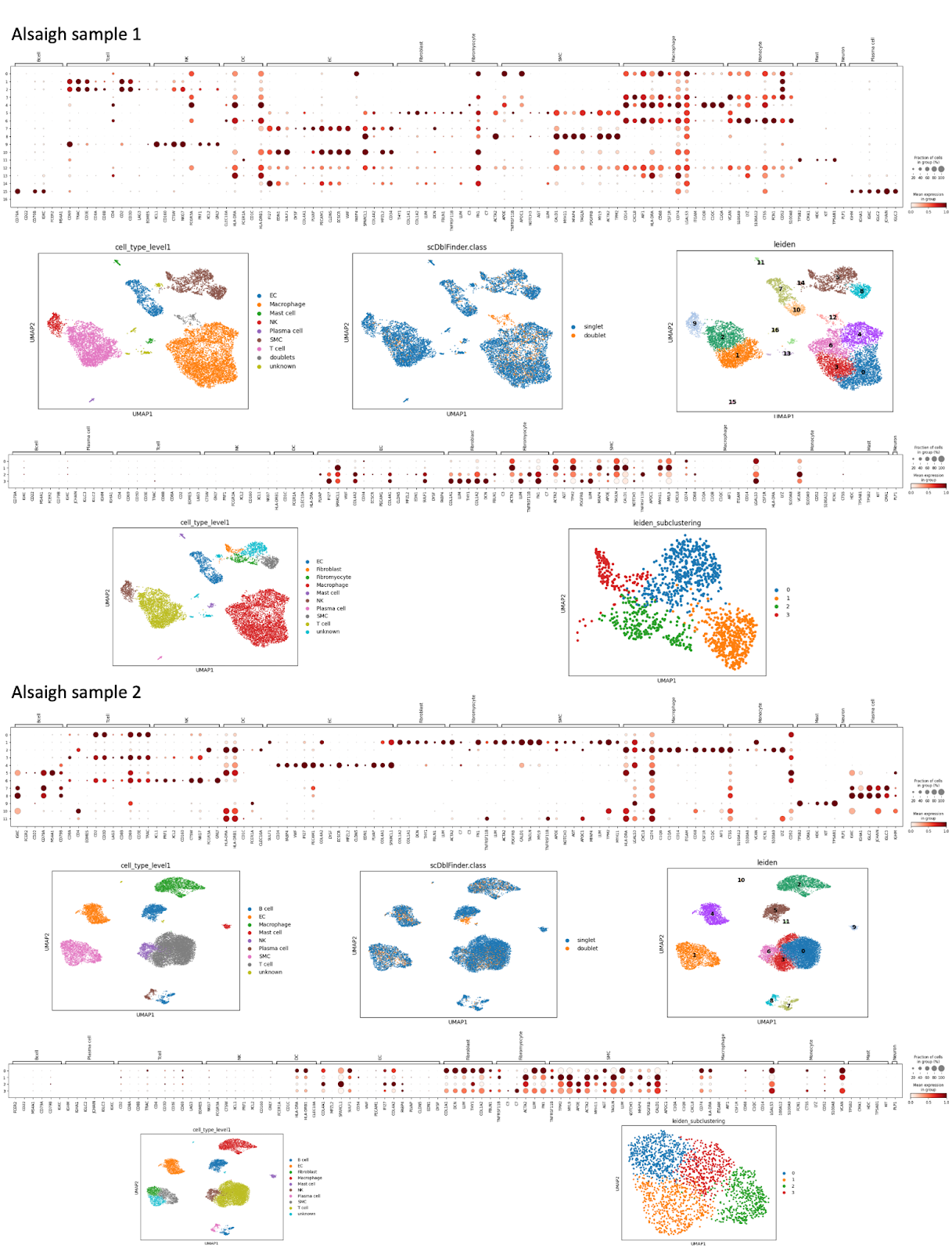

**Suppl. Figure 1***: Manual annotation of a subset of the samples in the atlas using marker genes.* For the manual annotation several samples were preprocessed and annotated with dedicated marker genes. For each sample the dot plot with the Leiden clusters on the y-axis and the marker genes on the x-axis is shown, together with the corresponding UMAPs of the annotated cells with Level 1, the doublet classification from ScDblFinder, and the Leiden clusters. In some cases, sub-clustering on specific clusters were performed and the corresponding dot plot, cell type UMAP and Leiden UMAP shown.

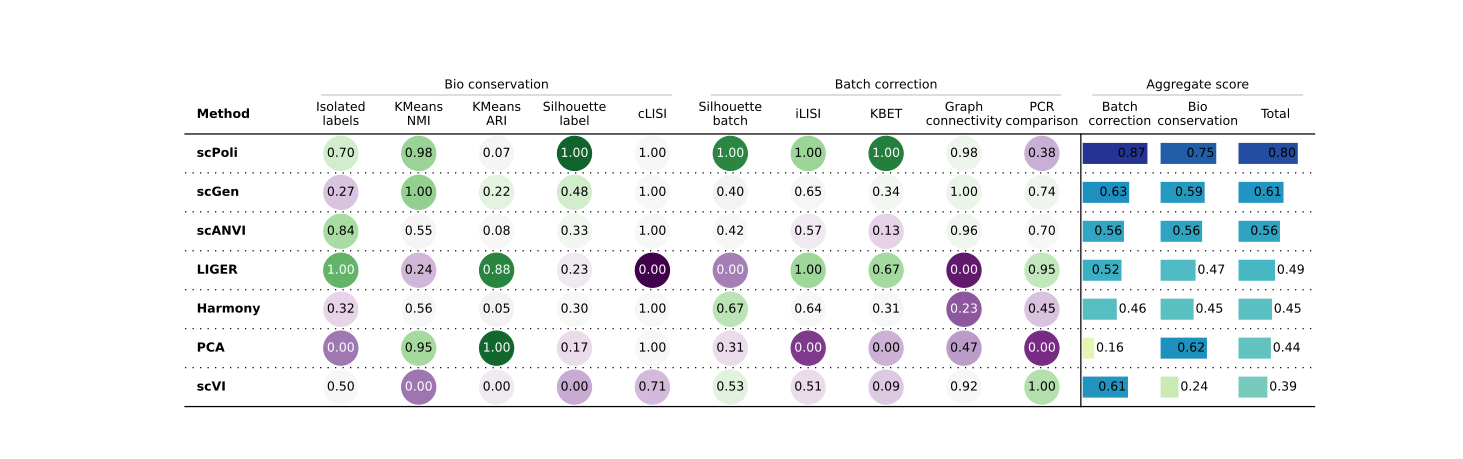

**Suppl. Figure 2:** *Benchmark of integration methods with scib-metrics on a subset of manually annotated samples.* For each method, ten metrics are calculated and normalized between 0 and 1 across the methods. Five Bio conservation and five batch correction metrics are aggregated in one total score. scPoli was the best performing method in bio conservation and in batch correction.

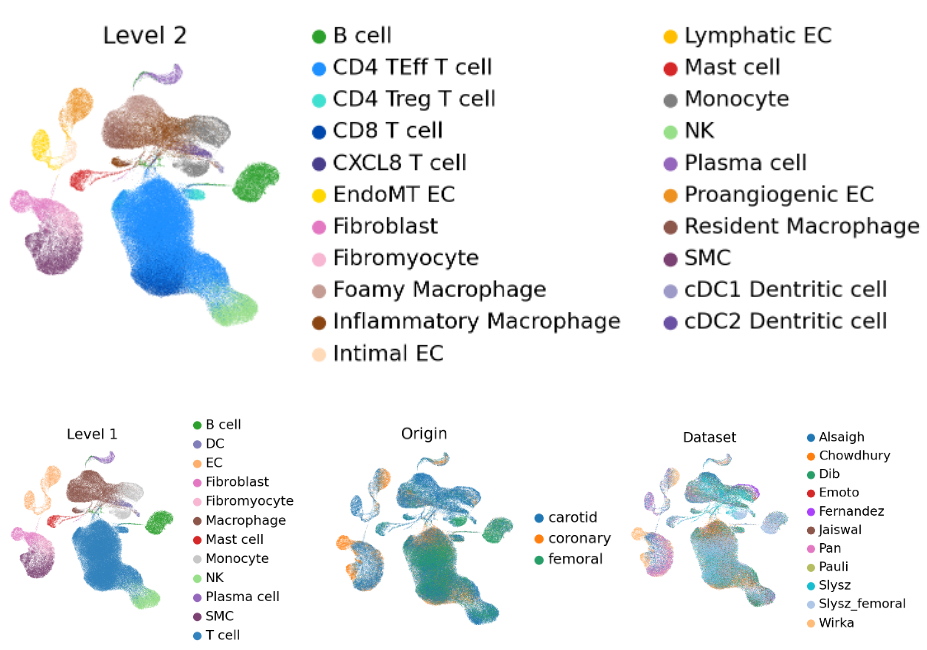

d

c

a

b

**Suppl. Figure 3***: Reference atlas after level 2 integration.* a) UMAP of the embedding after integration including Level 2 annotations. b) UMAP including the Level 1 annotations inferred from Level 2. c) UMAP with the origin site of the cells tissue. d) UMAP with the different datasets used in the reference atlas.

b

d

c

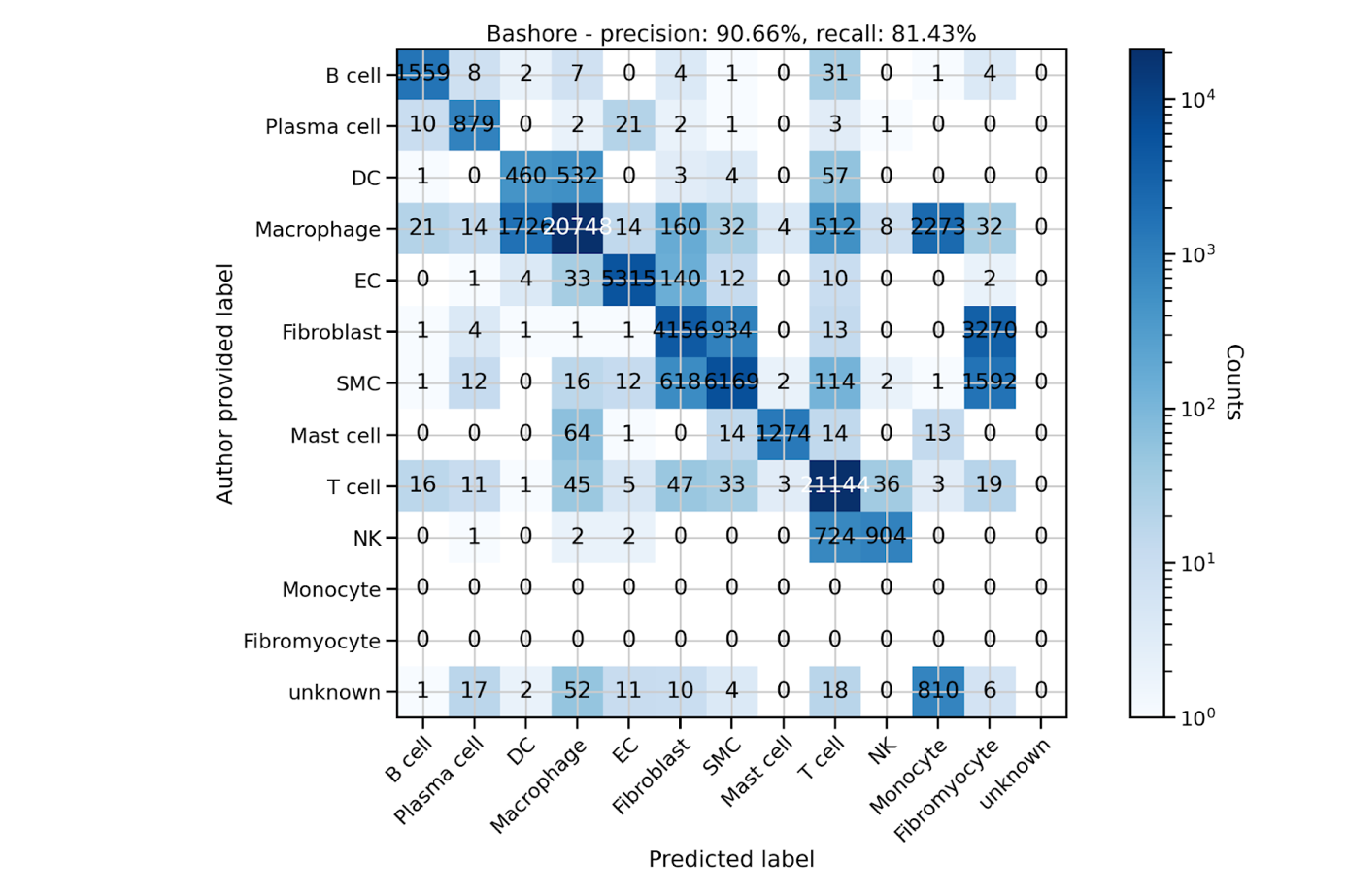

**Suppl. Figure 4:** *Validation of the cell type label transfer.* Confusion matrix of the Bashore et al. mappings with all cell types, including the cell types that are not shared between the author provided labels and our predicted labels. To compensate for the differences in cell type abundance the colormap is logarithmized and the overall precision and recall are weighted according to their abundance.

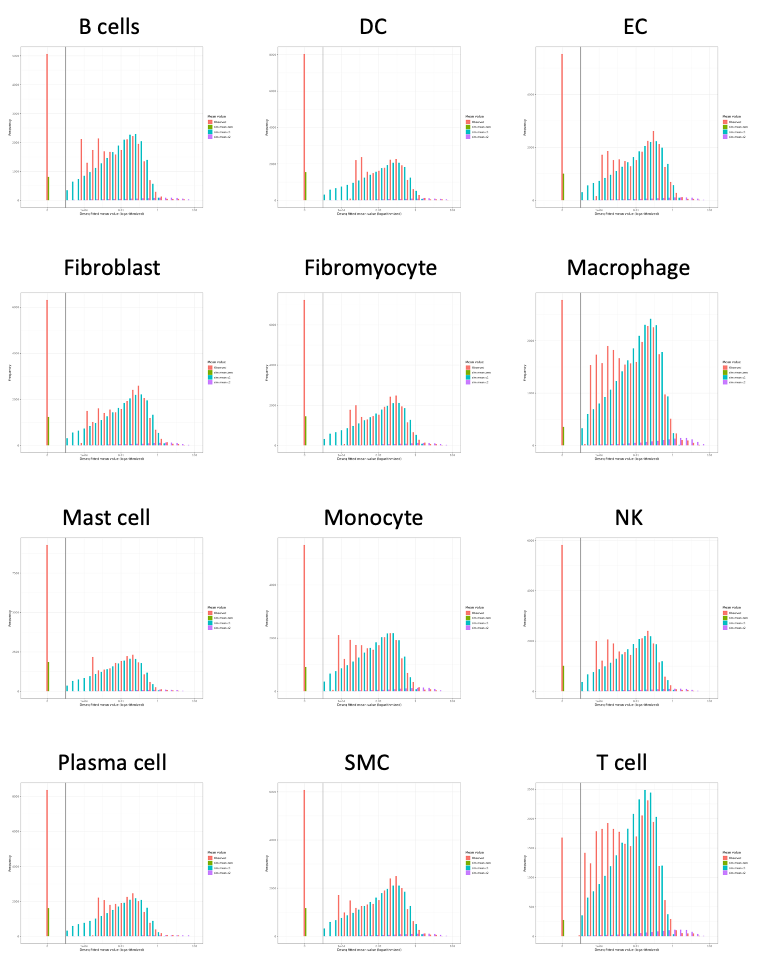

**Suppl. Figure 5***: Gene expression gamma prior fits used for scPower for each cell type using the atlas as a reference.* The panels depict the frequency of the Deseq fitted mean values (logarithmized) stratified in the zero component, two gamma distributions and the observed values per cell type.

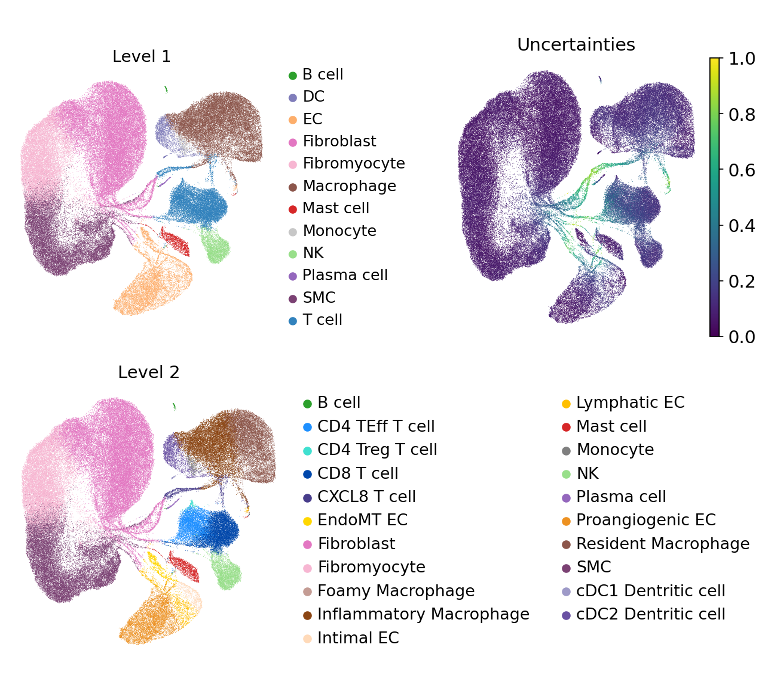

b

c

a

**Suppl. Figure 6***: Reference mapping of healthy arteries of the Hu et al. dataset on the plaque atlas.* UMAPs of the mapped Hu et al. cells to the plaque atlas depicted with Level 1 (a), Level 2 (c) and uncertainties of the model (b).

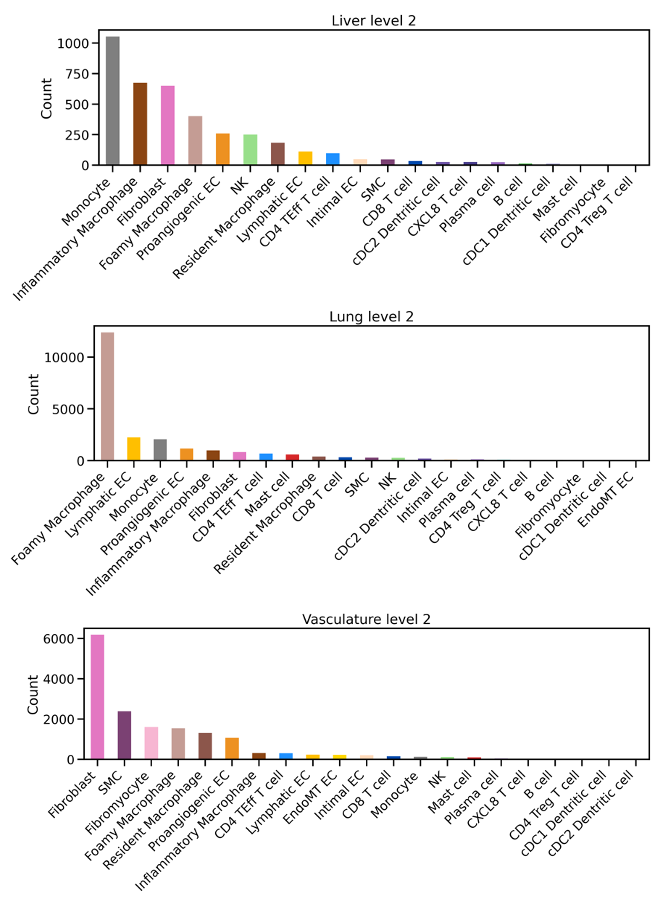

a

c

b

**Suppl. Figure 7***: Abundance of the predicted level 2 cell types in Tabula Sapiens.* Level 2 cell type abundance of the Tabula Sapiens subsets after mapping the Liver (a), Lung (b) and Vasculature (c) dataset to the plaque atlas.

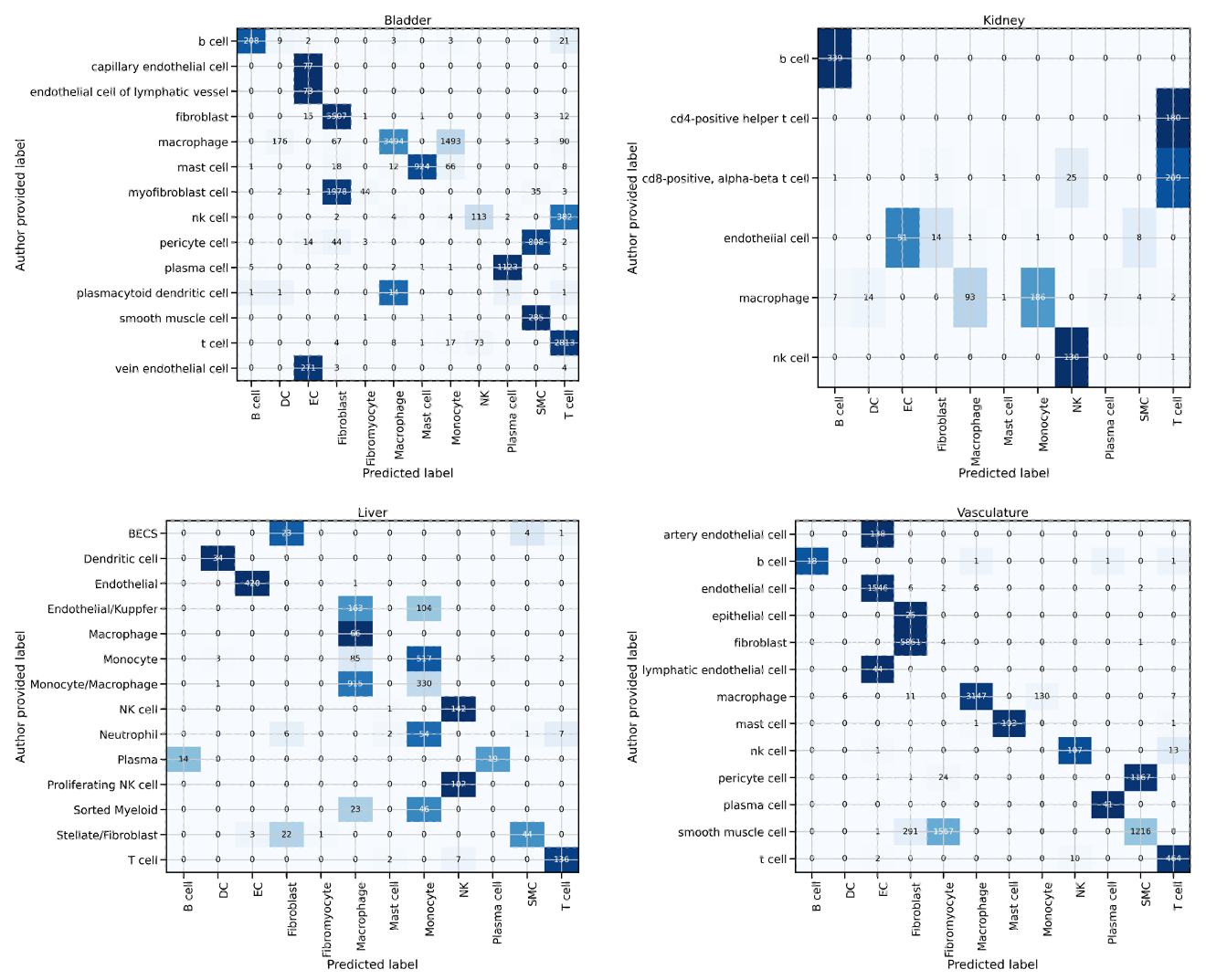

d

c

b

a

**Suppl. Figure 8***: Confusion matrices for the Tabula sapiens mappings.* Panels a-d show the confusion matrices for different data sets (data set name given in the panel headers) providing the number of cells with specific combinations of expert annotations (“Expert provided label” on the y-axis) and labels predicted by the atlas (“Predicted Label” on the x-axis). Organ specific cell types and cells with a higher uncertainty than 0.7 are removed. The colormap is normalized for each row to compensate for differences in cell type abundance.

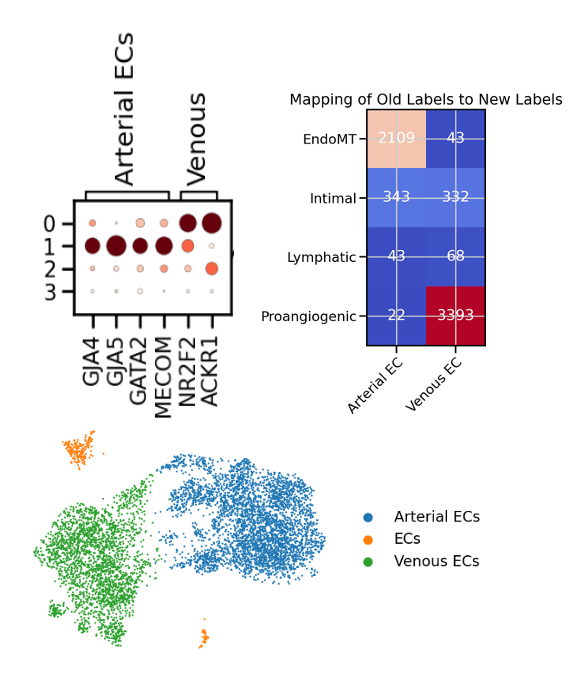

c

b

a

**Suppl. Figure 9***: Mapping of Reannotation of EC subtypes for deconvolution.* a) The Dot plot of the marker genes on the x-axis, and the Leiden clusters on the y-axis. b) depicts the mapping between the predicted labels of the atlas on the y-axis and the newly assigned labels on the x-axis. c) shows the UMAP of ECs with the new Labels assigned to the Leiden clusters.
